# Reconstructing the biogeography of a hunter-gatherer planet using machine-learning

**DOI:** 10.1101/2021.08.21.457222

**Authors:** Marcus J. Hamilton, Robert S. Walker, Briggs Buchanan, Damian E. Blasi, Claire L. Bowern

## Abstract

Estimating the total human population size (i.e., abundance) of the preagricultural planet is important for setting the baseline expectations for human-environment interactions if all energy and material requirements to support growth, maintenance, and well-being were foraged from local environments. However, demographic parameters and biogeographic distributions do not preserve directly in the archaeological record. Rather than attempting to estimate human abundance at some specific time in the past, a principled approach to making inferences at this scale is to ask what the human demography and biogeography of a hypothetical planet Earth would look like if populated by ethnographic hunter-gatherer societies. Given ethnographic hunter-gatherer societies likely include the largest, densest, and most complex foraging societies to have existed, we suggest population inferences drawn from this sample provide an upper bound to demographic estimates in prehistory. Our goal in this paper is to produce principled estimates of hunter-gatherer abundance, diversity, and biogeography. To do this we trained an extreme gradient boosting algorithm (XGBoost) to learn ethnographic hunter-gatherer population densities from a large matrix of climatic, environmental, and geographic data. We used the predictions generated by this model to reconstruct the hunter-gatherer biogeography of the rest of the planet. We find the human abundance of this world to be 6.1±2 million with an ethnolinguistic diversity of 8,330±2,770 populations, most of whom would have lived near coasts and in the tropics.

**Significance Statement:** Understanding the abundance of humans on planet Earth prior to the development of agriculture and the industrialized world is essential to understanding human population growth. However, the problem is that these features of human populations in the past are unknown and so must be estimated from data. We developed a machine learning approach that uses ethnographic and environmental data to reconstruct the demography and biogeography of planet Earth if populated by hunter-gatherers. Such a world would house about 6 million people divided into about 8,330 populations with a particular concentration in the tropics and along coasts.

---

In 2021, the global abundance of the human species is almost 8 billion. In 1921 it was about 2 billion, in 1821 1 billion, in 1621 about 500 million, and in 21 AD about 200 million (1). Prior to the emergence of agriculture in the Fertile Crescent about 10,000 BC, global abundance was likely no more than a few million (1). If human population growth saturates at 11 billion around 2100 AD as predicted by the UN (2, 3) human abundance will have increased 5000-fold in 13,000 years (Figure 1). However, reliable demographic statistics are available only for the last couple of centuries (4). Even with government funding and resources, national-scale population estimates in the 21st century cost billions of dollars and years of preparation (5). Reconstructing global scale phenomena in the deeper past is even more challenging as demographic parameters do not preserve in the archaeological or geological records (6). Moreover, the human species has a dynamic evolutionary history of adaptive radiation, range expansion, founder effects, bottlenecks, fluctuating climates, and cultural evolution building complex spatiotemporal dependencies into any prehistoric demographic estimates.

**Fig. 1.**
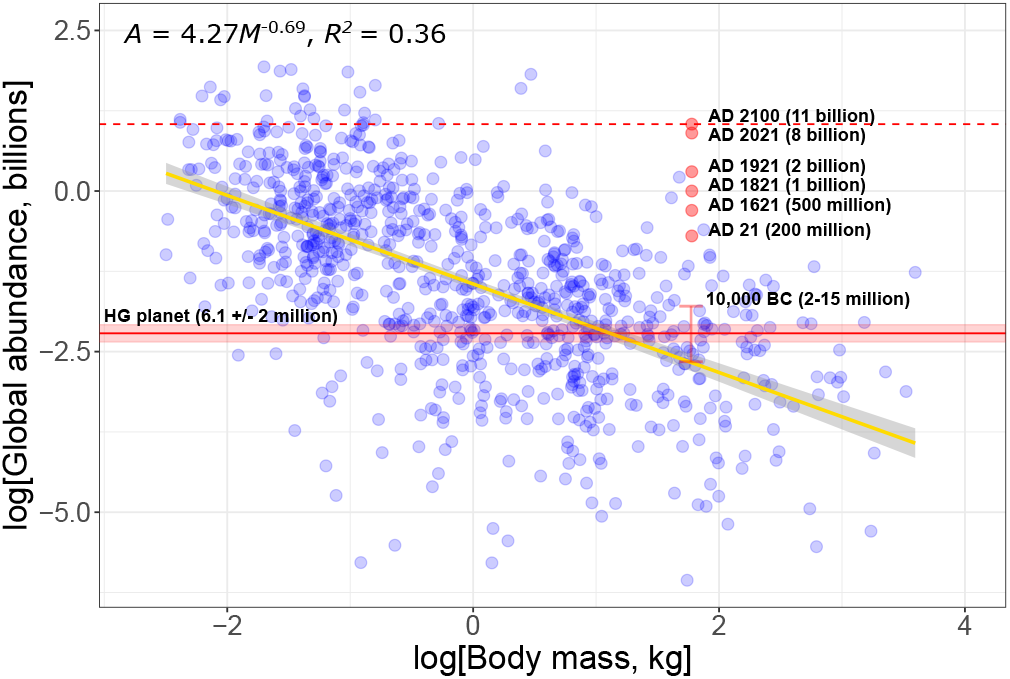
The total abundance of mammal species (blue data) and humans over time (red data) by body size on logged axes. The orange line is an OLS regression model fit to the mammal data, and the equation is top left. The dashed line is the United Nations prediction of 11 billion by AD 2100 and the solid line is the 6.1±2 million estimate we develop here. Human abundance will have increased 5,000-fold over the Holocene.

A reasonable zeroth-order anthropological model of global hunter-gatherer abundance is simply to multiply the average population density of recently-recorded ethnographic hunter-gatherers, 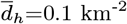, by the available terrestrial landmass of the planet, *A_h_* = 149 million km^2^, in which case global abundance would be 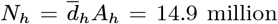. Similarly, a reasonable ecological estimate comes from the allometric scaling of mammal body size, *M_m_*, and species abundance, *N_m_* (Figure 1), where 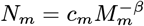, and so using the parameters from the fitted model in Figure 1, the predicted total abundance of a 60 kg mammal is 2.1 million. This range of 2 - 15 million bounds the range of estimates of human preagricultural abundance commonly found in the anthropological literature (1, 7–9). However, these estimates are often quite general in nature and rarely address the complex interactions that constrain hunter-gatherer biogeography and spatial ecology explicitly.

Given the complexity of human evolutionary history, a principled approach to developing more rigorous estimates of demography in the deep past is to consider a hypothetical world populated by ethnographic societies to serve as a null model. The spatial ecology of ethnographic hunter-gatherers at a coarse-grained scale is now well understood empirically and theoretically (8, 10–12). Because all energy requirements for maintenance, growth, and motility are derived from local ecosystems, environmental productivity constrains the diversity of hunter-gatherer adaptations in predictable ways (13, 14). For example, global variation in hunter-gatherer landscape use (11, 15), population density (11), and mobility (16–19) are all predicted by variation in environmental productivity. In this paper we use these insights to develop a machine-learning approach to predict the abundance, diversity, and biogeography of a hypothetical hunter-gatherer planet. In specific, we use extreme gradient boosting (XGBoost) (20), an advanced regression technique that uses a gradient boosting decision-tree modeling framework to build complex ensemble models of independent data to predict a dependent variable (see **Supplementary Information** for details).

## Model development

Our machine-learning approach begins by modeling hunter-gatherer population density as a function of a large matrix of biotic and abiotic environmental data comprised of 46 predictor variables. These data include climatic, biological, and geographic variables designed to capture environmental complexity. Given the results of this model, we then project predicted hunter-gatherer population densities across the planet based on the same matrix of environmental data. From these projections we establish estimates of the global abundance and diversity of hunter-gatherer populations. We then extract information from this reconstructed planet to examine the biogeographic distribution of population densities in terms of both the abiotic and biotic environments. For a detailed description of the data and modeling approach see the **Supplementary Information** attached to this paper.

### Modeling ethnographic data

A key question in comparative anthropology is how best to use ethnographic data to reconstruct scales of demography beyond those directly observed in contemporary populations (21, 22). The only direct source of information about hunter-gatherer lifestyles comes from observations of ethnographic societies (10), all of whom have been impacted by colonialization and the expansion of the industrialized world. Simple ethnographic analogies that equate case studies in the past with case studies in the present are problematic as analogies are not homologies, and so lack mechanistic equivalence. A more principled approach is statistical: Ethnographic data can be used to make inferences about parameters of interest and to measure the errors with which those inferences are made.

Although the ethnographic record of the Anthropocene is a non-random sample, most sources of bias are likely to inflate the range of variation we observe in the ethnographic present versus the past. For example, while the spread of diseases and environmental marginalization catastrophically reduced population numbers over the colonial era, accumulating technological innovations, the warm stable climates of the Holocene, and widespread interactions with non-foraging peoples likely increased populations over the long term. Thus ecological parameters in the present are likely *less* predictable solely from environmental data than in the past because ethnographically there are additional sources of variation.

Consider population density for example. Population size per unit area is correlated with environmental productivity as the energy density of landscapes fundamentally constrains the amount of biomass that can be supported per unit area (11). However, all populations naturally cycle through periods of growth, stability and collapse and so cannot be assumed to be in equilibrium (23, 24). Moreover, in many ethnographic hunter-gatherer societies energy budgets are supplemented using technologies not available in the deeper past including metals, firearms, and combustion engines, and lifestyles incorporating horticulture, agriculture, or market economies. However, there is no *a priori* reason to believe that these interactions break the fundamental ecological symmetry between space and energy, especially as the predicted correlations are observed in data (8, 11). As such, uniformitarian assumptions of ecological and demographic equivalence in the past and present can be considered uniformitarian principles (25, 26).

## Results

### XGBoost model of population density

The XGBoost model of hunter-gatherer population density returns a cross-validated *r*^2^ = 0.96 and so variation in hunter-gatherer population densities are highly predictable from environmental, climatic, and geographic variables. Figure 3 shows the top 29 predictors of most overall importance where SHAP values (Shapley additive explanations) are the marginal impact of each predictor variable on the model conditional on all other possible interactions. Three of the top four predictors show are minimum temperature constraints during the coldest and driest parts of the year indicating that annual climatic extremes play important roles. The second highest predictor is bird diversity, which likely correlates with broader environmental variation. The fifth variable is proximity to rivers, and the sixth variable is pathogen load, consistent with Tallavaara et al. (8). This is followed by several measures of landscape complexity, such as proximity to coastlines, rivers, marine resources, and water sources. Total annual precipitation is ranked 10th in overall importance in the model. Interestingly, terrestrial mammal diversity is ranked 22nd on the list of importance, likely because much of the predictive power is captured by higher-ranked environmental predictor variables. A strength of the XGBoost model over more traditional linear models is it’s ability to capture the complex nonlinear responses of predictor variables and their interactions (Figure 4).

Figure 5 shows the probability distribution of modeled population densities covers the same range as the observed ethnographic sample, but the distributions change shape. The average population density of the hypothetical world is 0.03±0.01 km^−2^ (Figure 5A) whereas the average population density of the ethnographic sample is 0.11 ± 0.07 km^−2^. The highest population densities observed ethnographically are much rarer in the reconstructed world and so the mean population density is considerably less. Thus the XGBoost model is capturing - and correcting - the over-sampling of high density ethnographic societies in the training data, such as those of the Pacific Rim of North America, and mutualists of tropical regions of Southern Asia and Central Africa, such as the Chenchu of India, the Agta of the Philippines, and the Bayaka of the Congo.

### Reconstructing the hunter-gatherer planet

Figure 2A shows the biogeographic distribution of predicted hunter-gatherer population densities in the reconstructed world, Figure 2B shows the global distribution of above-ground net primary production, and Figure 2C shows the global distribution of marine mammal diversity. The spatial distribution of population densities correlates with environmental productivity, with particularly high densities in the tropics (±26° N/S of the equator), and locally in other highly productive environments, including the southern Japanese archipelago, the east coast of Australia, and the temperate west coast of South America. Population densities are also high along coasts including the North American Pacific Rim, central America, much of coastal South America, Atlantic Europe, the Mediterranean Basin, the northern Persian Gulf and the south coast of the Caspian, coastal sub-Saharan Africa, southeast Asia, and much of coastal Australia. Population densities are often particularly high along the most diverse coasts.

**Fig. 2.**
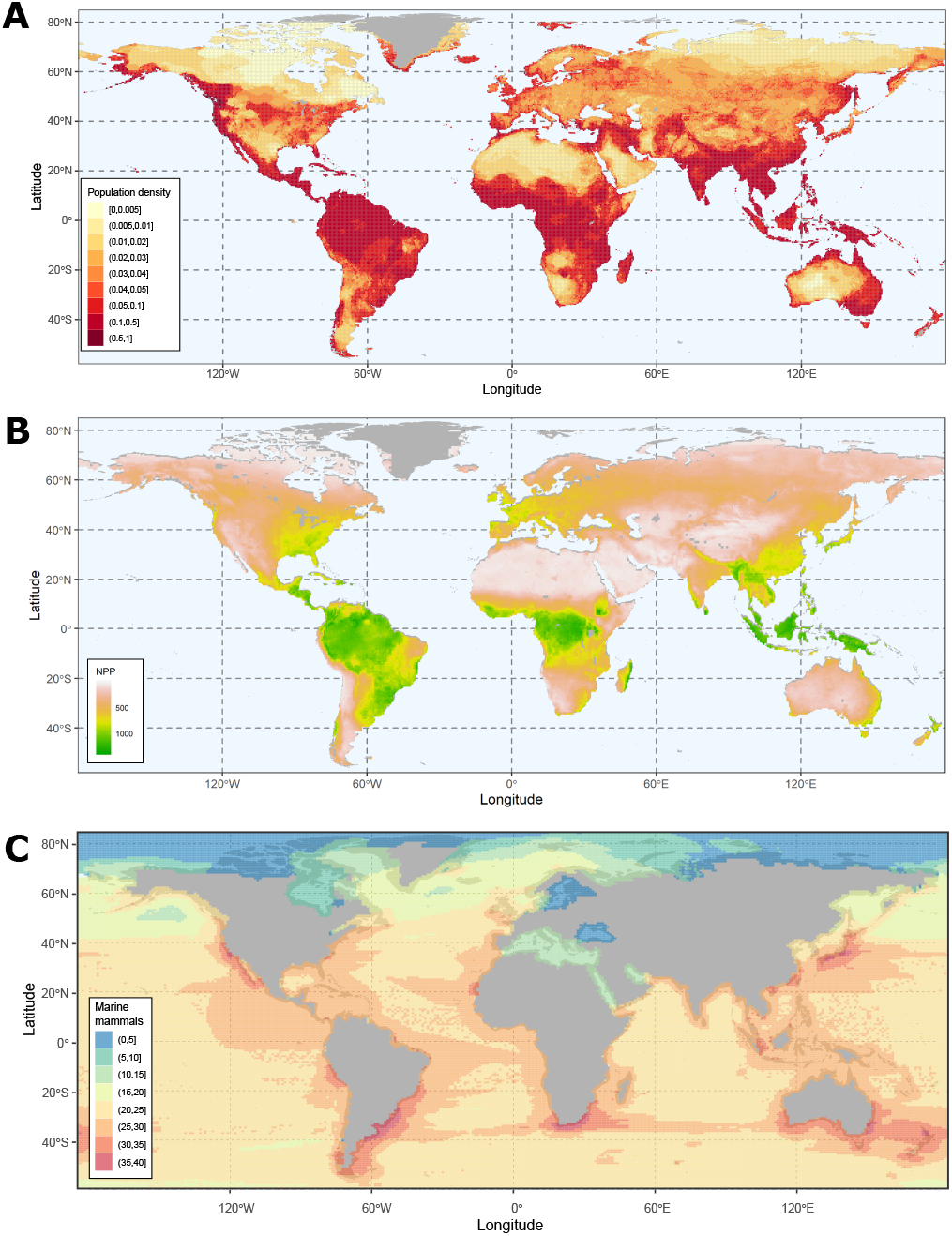
The global distribution of the reconstructed hunter-gatherer population densities (A), net primary production (B), and the biogeographic distribution of marine mammal diversity (C). Hunter-gatherer population densities tend to be highest in the tropics and in other regions of high net primary production, but also along many coastlines, particularly those with high marine mammal diversity. Outside the tropics and away from coasts population densities are much lower.

**Fig. 3.**
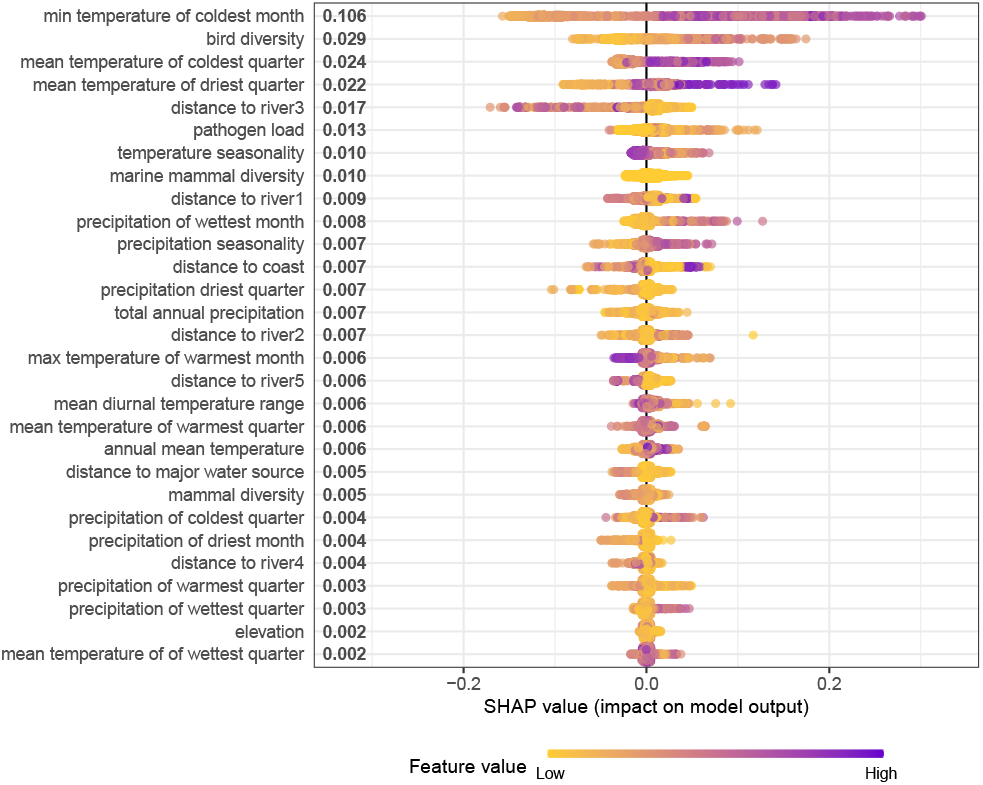
The marginal importance - SHAP values - of the predictors of hunter-gatherer population densities in the XGBoost model. Positive values to the right of center have positive marginal effects on population density and purple reflects higher impact. Negative values to the left and have negative effects and yellow reflect lower impact For example, high mean annual temperatures (ranked 4th) have a strongly positive impact on population densities and low values have a weakly negative impact. Less temperature seasonality has a high impact (ranked 11th) on population density.

**Fig. 4.**
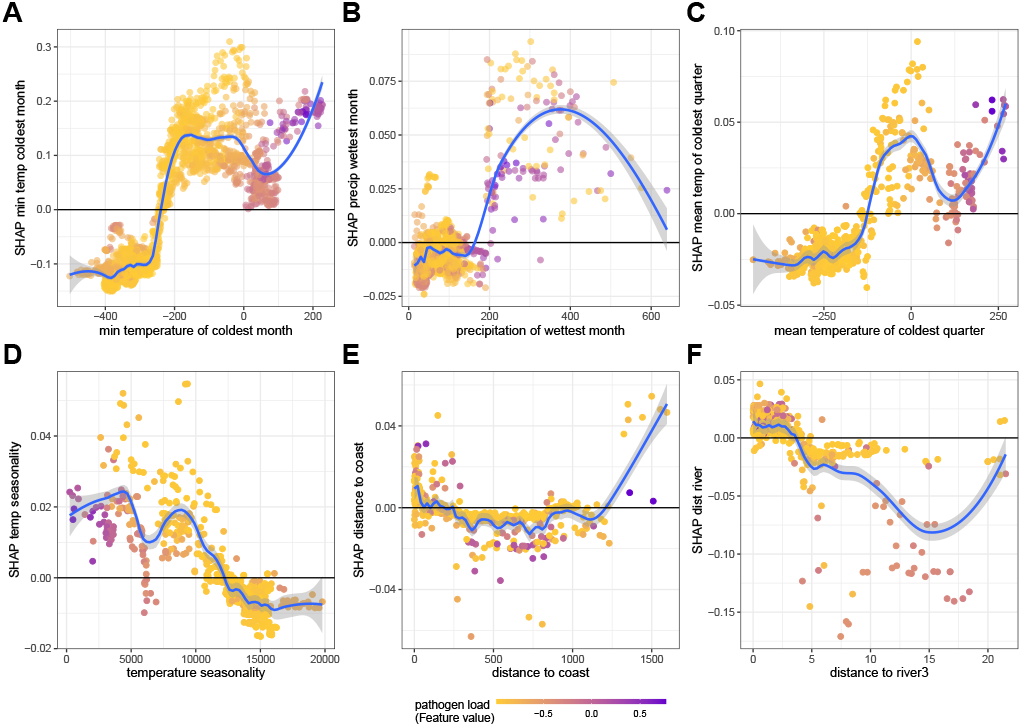
The range of nonlinear dependencies of the marginal impact of six predictor variables on hunter-gatherer population densities captured by the XGBoost model, color-coded by pathogen load. In each panel, the y-axis is the marginal dependency of the x-axis on population density, color-coded by pathogen load (yellow to purple). A-C) Temperature and precipitation variables have negative impacts on population density at low values, and positive at high, but the responses are highly nonlinear reflecting abrupt transitions in the responses. D) Increasingly seasonal temperatures have negative impacts on population density. E) The marginal effect of distance to coast is positive at low distances and negative beyond 300 km. However, at high distances (i.e., far inland) the effect becomes positive again, probably due to the proximity to other major bodies of water. F) The marginal effect of distance to tertiary branches of rivers decreases with distance and is negative after only a few kilometers.

**Fig. 5.**
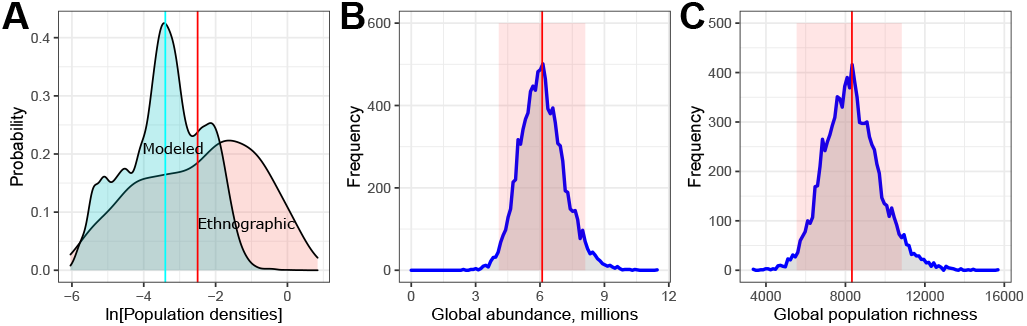
Population density, abundance, and diversity distributions from the XGBoost model. A) Modeled versus ethnographic population densities, where the vertical lines are the averages. B) The distribution of 10,000 Monte Carlo simulations of population abundance, with a mean of 6.1±2 million). C) The distribution of 10,000 Monte Carlo estimates of ethnolinguistic population richness, with a mean of about 8,330±2,770).

### Abundance and diversity estimates

To estimate the expected abundance and diversity of a hunter-gatherer planet we ran 10,000 Monte Carlo simulations of the XGBoost model, sampling with replacement from the ethnographic data. Each simulation produced an abundance estimate by summing over the global population density map it produced, creating the distributions in Figure 5B and 5C. The average total human population abundance was 6.1 (4.1-8.1) million. The confidence limits around this prediction, ±2 million, reflect the variance of the original ethnographic sample. Total abundances over 10 million were rare (9/10000 runs) and no single run exceeded 11.5 million. Similarly, total abundances of less than 3 million were rare (8/10000 runs) and the minimum abundance from a single run was 2.4 million.

To estimate ethnolinguistic diversity for each simulation we divided the total population abundance of each run by a bootstrapped geometric mean ethnolinguistic population size from the original ethnographic sample data. The boot-strapped geometric mean population size of the ethnographic societies was 732 (723-740). The average global ethnolinguistic population diversity was 8,330 (5,579–11,082). The minimum observed diversity was 3,166 and the maximum was 15,299.

Table 1 shows the predicted abundances and diversities across continents. Average population densities are predicted to have been highest in South America and Africa but considerably lower in Eurasia, Australia, and North America. The predicted number of ethnolinguistic populations in Australia, 370 130, is consistent with ethnographic estimates at European contact. While most of the Pacific was never colonized by hunter-gatherer societies, it is interesting that the low predicted diversity in Table 1 (140 ± 50) includes Papua New Guinea, ethnographically the most linguistically diverse region of the planet (27). However, the majority of societies on Papua New Guinea constituting this high diversity are horticulturalist-farmers not hunter-gatherers. Although hunter-gatherer population densities of Papua New Guinea are predicted to have been relatively high (Figure 2C), the model predicts relatively low diversity given the size of the island, suggesting that the high levels of diversity observed ethnographically are associated with non-foraging lifestyles.

**Table 1.**
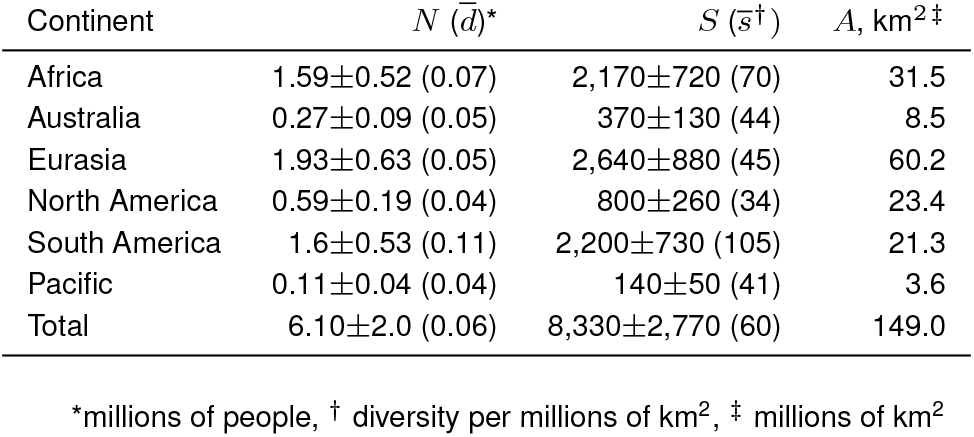
The distribution and errors (*x* ± 1.96*σ*) of reconstructed human population abundance, 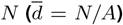, ethnolinguistic diversity, 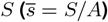, and landmass, *A*, across continents

### Biogeography and biodiversity estimates

Figure 6 shows the majority of hunter-gatherer populations (55%) would occur in tropical biomes, particularly evergreen forests and savannahs. Although pathogen loads are high in the tropics – and are an important predictor of population densities in the XGBoost model (see Figures 3 and 4) – tropical biomes are not only environmentally productive, but perhaps more importantly, there are many coasts in the tropics, especially in southern and southeast Asia, northern Australia, and Papua New Guinea.

**Fig. 6.**
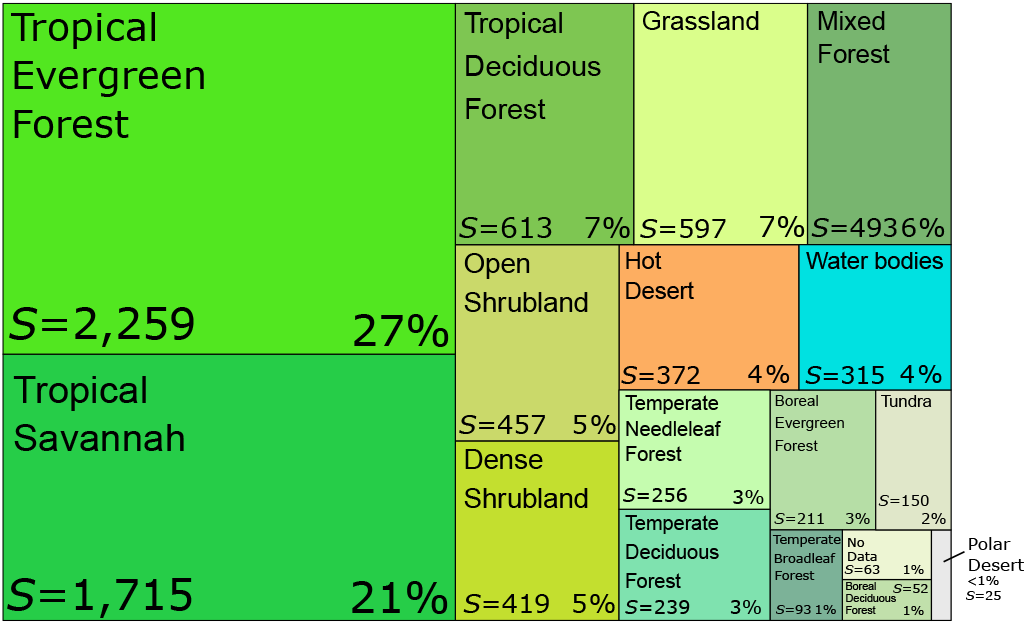
A tree map of the distribution hunter-gatherer cultural diversity (*S* and %*S*) distributed across the 16 biomes of the reconstructed hunter-gatherer planet, for a total diversity of 8,330. 55% of hunter-gatherer diversity occurs in tropical biomes followed by grasslands, mixed forests, shrublands, hot deserts, and islands. Less than 15% of cultural diversity occurs in temperate, boreal, tundra, and polar biomes.

Figure 7 shows the distributions of various measures of biodiversity and cultural diversity for five metrics of interest, where the blue line represents the availability of landmass along the gradient represented; data above the blue line occur at high densities (i.e., over-represented), and data below the line occur at low densities (i.e., under-represented). A striking result is that in all plots hunter-gatherer cultural diversity correlates closely with both mammal and bird species diversity. Figure 7A shows that all forms of diversity are over-represented in the tropics (±23.5° N/S) and under-represented at more northerly latitudes (>23.5° N). Figure 7B shows that hunter-gatherer diversity is over-represented in warm (> 20°C, Figure 7C), wet (>1000 mm^−yr^, Figure 7D), highly productive environments (>500 g C m^−2^, Figure 7B). Correspondingly, there is an under-representation in colder (<5°C, Figure 7C), drier (<700 mm^−yr^, Figure 7D), less productive environments (<500 g C m^−2^, Figure 7B). Moreover, Figure 7E shows that diversity decreases rapidly with distance from the coast where about 25% of global hunter-gatherer abundance occurs within 25 km of the coast, and 50% occurs within 150 km. Figure 7F shows that hunter-gatherer population densities (red high, blue low) are highest near the equator, particularly along coasts.

**Fig. 7.**
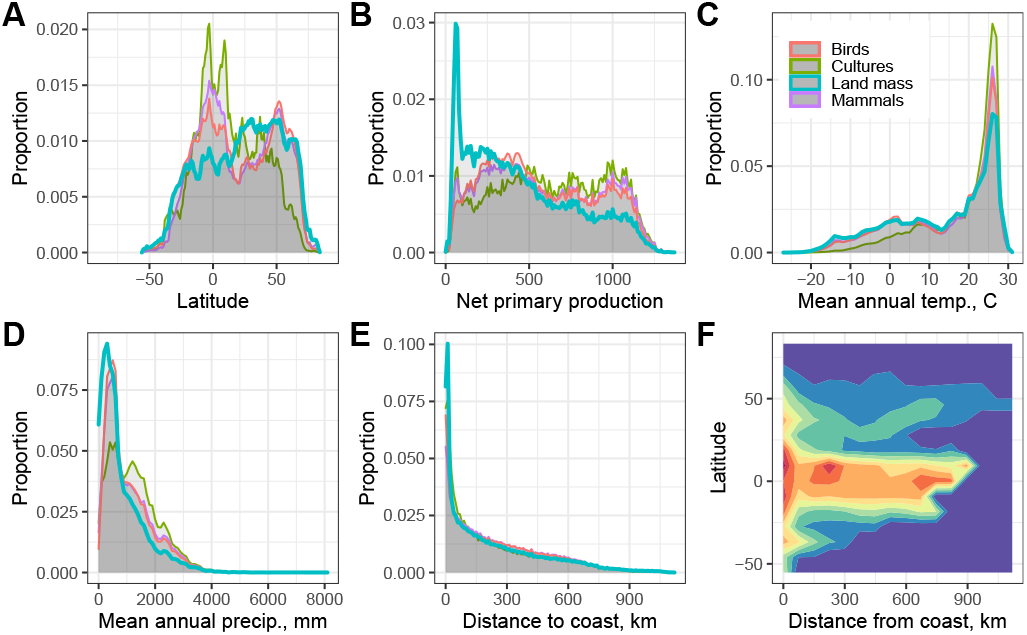
Distributions of biodiversity and cultural diversity by A) latitude, B) net primary production, C) mean annual temperature, D) mean annual precipitation, E) distance from coast, and F) population density (red high, *d* = 0.2, blue low, *d* = 0) as a function of latitude and distance from coast.

## Discussion

The hypothetical hunter-gatherer planet Earth we reconstruct here would have an abundance of 6.1±2 million people with a ethnolinguistic diversity of 8,330±2,770 populations. The majority would occur within tropical biomes (Figure 6) and one quarter of all hunter-gatherers would live less than one day’s walk from a coast (25 km). Figure 2C shows that marine mammal diversity is generally high along continental coasts at all latitudes below 60°N, and regions of high hunter-gatherer population density in environments with low net primary production often coincide with particularly productive coasts, including southern South America, the south and east coasts of Africa, coastal south Asia, coastal Australia, and the west coast of North America (Figures 2A and 2C). Marine mammal diversity is high throughout the tropics, and coupled with highly productive terrestrial and marine environments (see Figures S3), these biomes support the most diverse and dense hunter-gatherer populations on the planet. Outside the tropics and inland from coasts hunter-gatherer population density drops rapidly: Figure 6 shows only 15% of cultural diversity (thus abundance) occurs in temperate, boreal, tundra, and polar environments combined, which constitute 40% of the planet’s terrestrial surface area. Our results suggest that the ethnographic record of terrestrial, low density societies is biased not necessarily due to marginal environments, but because most are not coastal. Importantly, Figure 5A shows that the importance of coasts in the model is not a statistical artifact of the over-sampling of dense, complex, sedentary societies of the North American Pacific Rim, or tropical forager-farmer mutualisms in the ethnographic record, for example: these kinds of high density societies are in fact exceedingly rare in the reconstructed world, where average global population densities are substantially lower than in the ethnographic data.

While diversity decreases rapidly with the distance from coasts, Figure 7E shows there is no evidence for positive selection for coasts as the decline in diversity simply maps onto the geometric decrease in land availability with increasing distance inland. Thus hunter-gatherer population density (and biodiversity) follows an ideal free distribution with respect to landmass (28). In contrast, Figure 7B shows highly productive terrestrial environments are particularly dense with biological and cultural diversity and much lower in low productivity environments. This finding is consistent with well-established principles of biogeography where both biological and cultural diversity are higher in the tropics than predicted by environmental productivity and land mass alone (29, 30). The classic image of low density, highly mobile hunter-gatherers moving over vast distances may be a relatively rare adaptation to specific landscapes with low environmental productivity and no direct access to coasts.

Low productivity environments far inland would have been more common in the cooler, drier climates of the late Pleistocene, as sea levels were considerably lower increasing land-mass availability and therefore the average distance to the coast. Cooler, drier climates reduced the elevation range of biomes and thus reduced the spatial extent of the tropical zone globally (31). Under these conditions low population density adaptations would have been more common than we see ethnographically, but the majority of populations would still have occurred adjacent to coasts. In fact rich, diverse coastal environments may have been even more important given the decreased availability or terrestrial resources. If 25% of hunter-gatherer populations lived within 25km of the coast in the late Pleistocene, sea level rise over the Holocene would have had a dramatic impact on the archaeological visibility of late Pleistocene hunter-gatherer lifestyles, especially in areas of shallow continental shelf (32), such as Beringia (33), Dogger Bank (34), or the Sunda shelf (35), adding to the archaeological over-representation of low density lifestyles.

Importantly, our reconstructed world of ethnographic hunter-gatherers in the Anthropocene is not the pre-agricultural world of stone age hunter-gatherers. The Holocene is a particularly warm, wet, and stable period of recent Earth history (36), suggesting average population densities would be considerably lower in the late Pleistocene. Moreover, ethnographic societies are neither a cultural nor technological analogue of past foraging societies. All ethnographic societies have been impacted by the global expansion of industrialized societies over the last 500 years. Widespread outbreaks of disease, violence, forced relocation, ecological marginalization, and habitat loss led to the decimation of indigenous societies in many parts of the world, and many ethnographic societies were first encountered over this period. All ethnographic hunter-gatherer societies have widespread interactions with non-foraging societies of the Anthropocene, incorporating innovations such as medicines, clothing, tools, metals, firearms, and combustion engines whenever useful. No ethnographic societies now use stone tools, a central feature of foraging socioeconomies throughout human evolutionary history. Many contemporary foragers practice horticulture, others participate in long-term mutualisms with farming societies, and many participate in market economies, all of which are post-Pleistocene adaptations. Similarly, the sociopolitical diversity of hunter-gatherers in the late Holocene/Anthropocene undoubtedly includes societies with institutionalized economic and political asymmetries that were likely much rarer, if not absent, in preceding time periods in many parts of the world. Particularly important here is that many of these Holocene adaptations included the increased specialization of marine coastal environments.

The relevant anthropological question here is understanding how these historical, cultural, and technological contingencies may have impacted the range of population densities we see in the ethnographic record compared to those in the past. Ethnographically there is a thousand-fold variation in population densities, from 0.003 km^−2^ in the North American Arctic and the Western Desert of Australia to 3 km^−2^ along the Californian coast, with a sample average of 0.11 km^−2^. It is reasonable to assume hunter-gatherer population densities have always been particularly low in warm and cold deserts, likely at the scales we observe ethnographically, but the maximum densities, such as those found in the sedentary societies of the North American Pacific Rim, are relatively recent. It is not known whether similarly dense populations occurred in the late Pleistocene, but if so they were likely much rarer than in the ethnographic record. As such, the range of population densities we observe ethnographically is probably wider than in prehistory, and if so the demographic parameters we estimate here are likely upper estimates of the pre-agricultural hunter-gatherer planet Earth.

## Conclusion

Our machine-learning approach provides a detailed reconstruction of the biogeographic distribution of hunter-gatherer societies. Populations would have been particularly dense and diverse along productive coasts and in tropical biomes. Assuming our model overestimates the population density of the prehistoric world, the pre-agricultural hunter-gatherer world likely never exceeded 6.1±2 million people. 6.1±2 million should also be seen as a principled estimate of the scale of the human species if all energy and materials were extracted from local environments in a counterfactual Anthropocene. Human abundance is now near 8 billion, and will be near 11 billion by the end of the century, in which case the evolution of the post-foraging world will have seen a 5,000-fold increase in human abundance over the Holocene.

## Materials and Methods

Our model uses extreme gradient boosting implemented through the *xgbTree* option in the *caret* package in R (37). For a complete description of the XGBoost algorithm see (20, 38). Extreme gradient boosting is an ensemble learning algorithm that uses a matrix of data to sequentially build a large number of regression tree models that maximize predictions of the dependent data by aiming to reduce model residuals in each iteration (see Figure S2). In our case the dependent variable is hunter-gatherer population density and the predictive data is a matrix of environmental data composed of 46 variables. The data include 17 climatic variables of temperature and precipitation, 3 variables of biodiversity data (mammal, bird, and marine mammal species diversity), 16 biome variables, 7 variables of physical landscape complexity, and pathogen data (see **Supplementary Information** for further discussion and sources). All data and R code are available in the **Supplementary Material** of this paper.

## ACKNOWLEDGMENTS

We appreciate comments on previous versions of this manuscript from J. Robbie Burger, Miikka Tallavaara and members of a working group on Human Macroecology funded by the Center for Biological Diversity at the University of Arizona in February, 2020, where this work was first presented and discussed.

